# Effect of external cues on clock-driven protection from Influenza A infection

**DOI:** 10.1101/2025.03.03.641279

**Authors:** Oindrila Paul, Thomas G. Brooks, Lora Assi, Alisha Shetty, Martina Towers, Kaitlyn M Forrest, Alecia Cameron, Amita Sehgal, Gregory R. Grant, Shaon Sengupta

## Abstract

Influenza and other respiratory viral pathogens are leading causes of mortality and morbidity. We previously demonstrated that circadian rhythms confer temporal protection from influenza infection. Here, we investigated whether this protection requires rhythmic function after the initial infection by manipulating environmental cycles. We demonstrate that disrupting environmental lighting cues within a critical window of vulnerability abrogates time-of-day specific protection. This poor outcome is mediated by a dysregulated immune response, evidenced by the accumulation of inflammatory monocytes and CD8^+^ cells in the lungs and a transcriptomic profile indicative of an exaggerated immune response. Disruption of the light cycle does not affect outcomes in a clock mutant, indicating that it acts by compromising endogenous timekeeping. Importantly, rhythmic meal timing mitigates the adverse effects of disrupted light cycles, suggesting that external cycling cues, which act through different body clocks, can substitute for each other. Our findings highlight the crucial interplay between environmental factors and endogenous clocks in shaping influenza outcomes, offering significant translational potential for improving the care of critically ill patients with respiratory viral infections.

## Introduction

Circadian rhythms provide an anticipatory protective mechanism in the face of environmental challenges, including the threat of infections. In mammals, the master pacemaker resides in the suprachiasmatic nucleus (SCN)^1^. However, almost every cell has its cell-intrinsic clock or molecular clock. Rhythmic external cues or zeitgebers synchronize central and peripheral clocks, ensuring optimal organismal function^2^. Light is the most important zeitgeber for the SCN, which then synchronizes other body clocks; however, other organs and cells are more sensitive to entrainment to organ-specific cues. Acute or chronic disruptions of entraining cues are well known to disrupt synchronization between the central and peripheral clocks, with their harmful effects best documented in chronic states such as shift work^3,4^. However, the impact of external entrainment in an acute context, mainly as it affects host defense, is less well understood.

Previously, we showed that the circadian clock confers time-of-day-specific protection from influenza infection. Mice infected at dawn had a threefold better survival rate than mice infected at dusk. This advantage was lost in mice with a genetically disrupted circadian clock *(Bmal1*^*-/-*^)^*5*^. However, mechanisms underlying this time-of-day effect remain unclear-for instance, is cycling physiology required only at the time of infection, or does it need to be sustained for a period post-infection? The extended time frame (8-10 days) from infection to peak mortality in the influenza model renders it well-suited to address this issue. This study examines how zeitgebers influence time-of-day specific protection in clock-intact hosts. We hypothesize that sustained external cues are necessary to maintain this protection after initial infection. Using light-dark cycles and meal timing as model zeitgebers, we assessed their impact on time-of-day variations in influenza infection outcomes.

### Time-of-day specific protection from influenza infection is lost after constant light exposure

We have previously shown that mice infected at dawn had threefold higher survival than mice infected at dusk^5^. We hypothesized that abrogation of cyclical photic cues caused by constant light exposure would abolish the time-of-day specific protection from IAV driven by the endogenous circadian clock. By circadian terminology, ZT0 refers to the time when the lights turn “ON,” and ZT12 is when the lights turn “OFF”; therefore, ZT23 refers to the time just before the lights go on or just before the onset of the rest phase, and ZT11 refers to the time just before the lights go off or before the onset of the active phase. As in our prior studies, C57Bl/6J adult mice were infected at either ZT23 or ZT11 with Influenza A Virus (IAV; H1N1; PR8-30 PFU) and maintained in 12 hr light-dark (LD) cycling conditions. A subset of mice infected at ZT23 were moved to constant light conditions on day 4 post-infection (p.i.), remaining there throughout the study [ZT23(LL-D4)] [Fig 1A]. Animals were weighed and monitored daily for 14 days.

**Fig 1.**
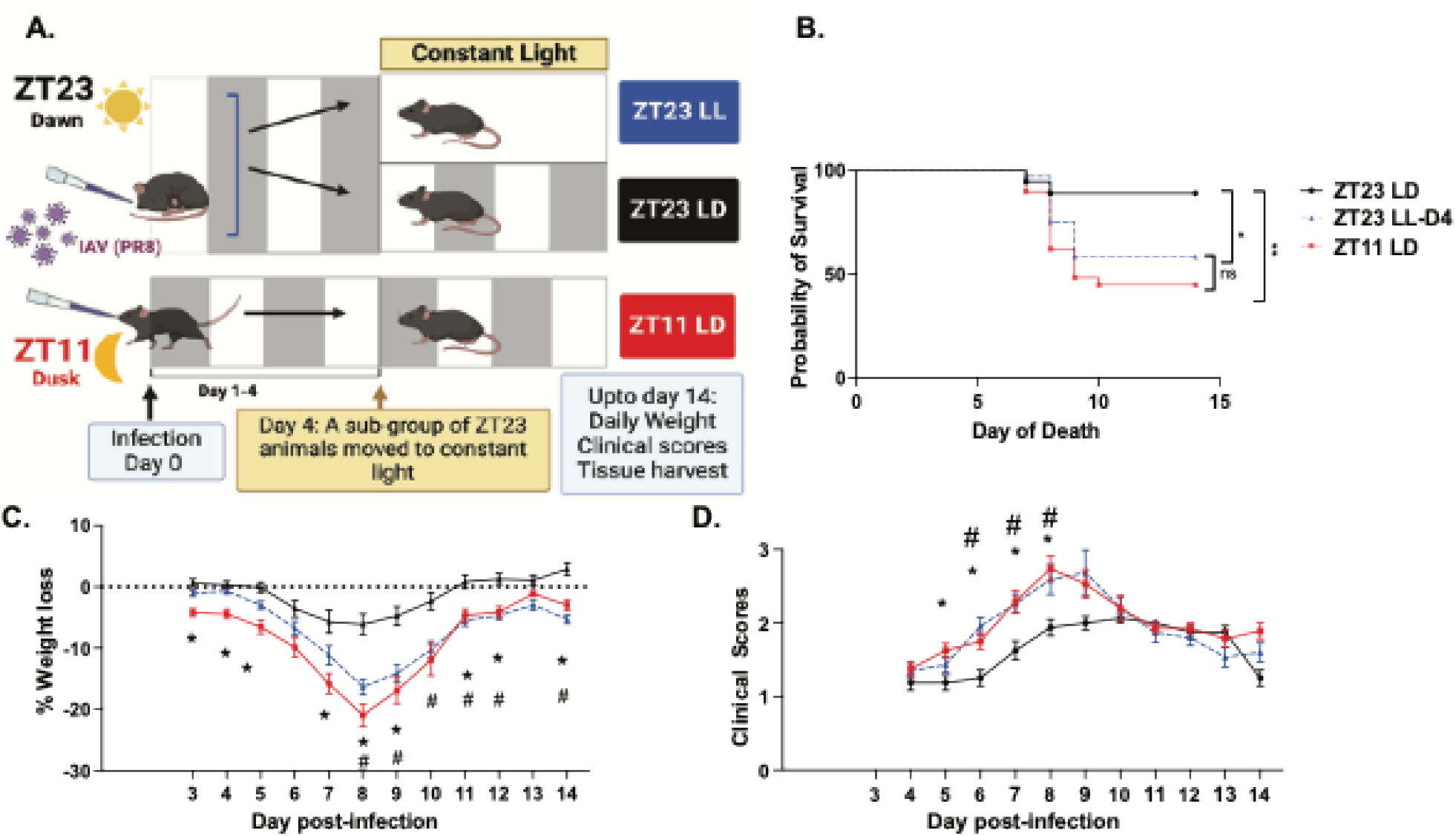
Constant light exposure abrogates the specific time of day protection following influenza A infection (A) Experimental model (B) Survival (n = 18–38 per group, **p<0.01 log-rank test, *p<0.05 log-rank test from three independent experiments) (C) Weight loss trajectory (n=18-38 per group, *p<0.001 ANOVA for repeated measures) (D) Average clinical score as a representation of disease progression. (n=18-38 per group, *p<0.001 ANOVA for repeated measures) following IAV infection. All data pooled from 3-5 independent experiments * = ZT23 vs ZT11; # = ZT23LD vs ZT23 LL; ns = non-significant

Consistent with our previous work, mice kept in 12 hr LD cycling and infected at ZT23 [ZT23(LD)] had significantly better survival [88.88% in ZT23(LD) vs 44.82% in ZT11(LD); p<0.01 by Mantel-Cox log-rank test] than the mice infected at ZT11(LD) [Fig 1B]. However, the ZT23(LL-D4) group had significantly lower survival [58.33% in ZT23(LL-D4) vs 88.88% in ZT23(LD); p < 0.05 by Mantel-Cox log-rank test] than the ZT23(LD) group [Fig 1B]. This was comparable to the mice infected at ZT11(LD) [44.82% survival in ZT11(LD) and 58.33% in ZT23(LL-D4)]. This effect is also reflected in the weight loss trajectory, where the ZT23 infected groups are identical until day 4 post-infection (p.i.) [Fig 1C]. However, thereafter, the weight loss is more pronounced in the ZT23(LL-D4) than the ZT23(LD) group, eventually becoming indistinguishable from the ZT11(LD) group [Fig 1C]. We also monitored these mice for sickness behavior using a previously validated scoring system [Supplementary Table 1] based on activity level, behavior, and respiratory distress^5,6^—the higher the score, the sicker the animal. Like the ZT11(LD) mice, the ZT23(LL-D4) group had consistently higher scores than the ZT23(LD) group, suggesting higher morbidity with constant light exposure [Fig 1D]. Together, these data suggest that disruption of light-dark cycling, even after the initial few days of influenza infection, abrogates the clock-driven time-of-day specific protection from influenza-induced mortality and morbidity.

### Constant light exposure does not affect ultimate clearance of virus

We quantified viral titers in lung tissue to determine whether environmental light disruption affects influenza A virus (IAV) clearance. Since the disruption of light cycling was introduced on day 4 post-infection (p.i.), viral titers were measured on days 6 and 8 p.i. to allow the host to experience the effects of this perturbation. For this dose of the IAV, we expect days 6 and 8 p.i to represent time points that follow peak viral titers and likely represent viral clearance. No significant difference in viral burden was observed between the ZT23(LD) and ZT23(LL-D4) groups on either day [Fig. 2A]. However, the ZT23(LD) group exhibited significantly lower viral titers than the ZT11(LD) group on day 6, consistent with our previous work. The ZT23(LL-D4) group has viral titers intermediate between the ZT23(LD) and ZT11(LD) groups. However, by day 8, all groups had cleared the virus. These findings suggest that light-dark cycle disruption does not significantly affect IAV clearance.

**Fig 2.**
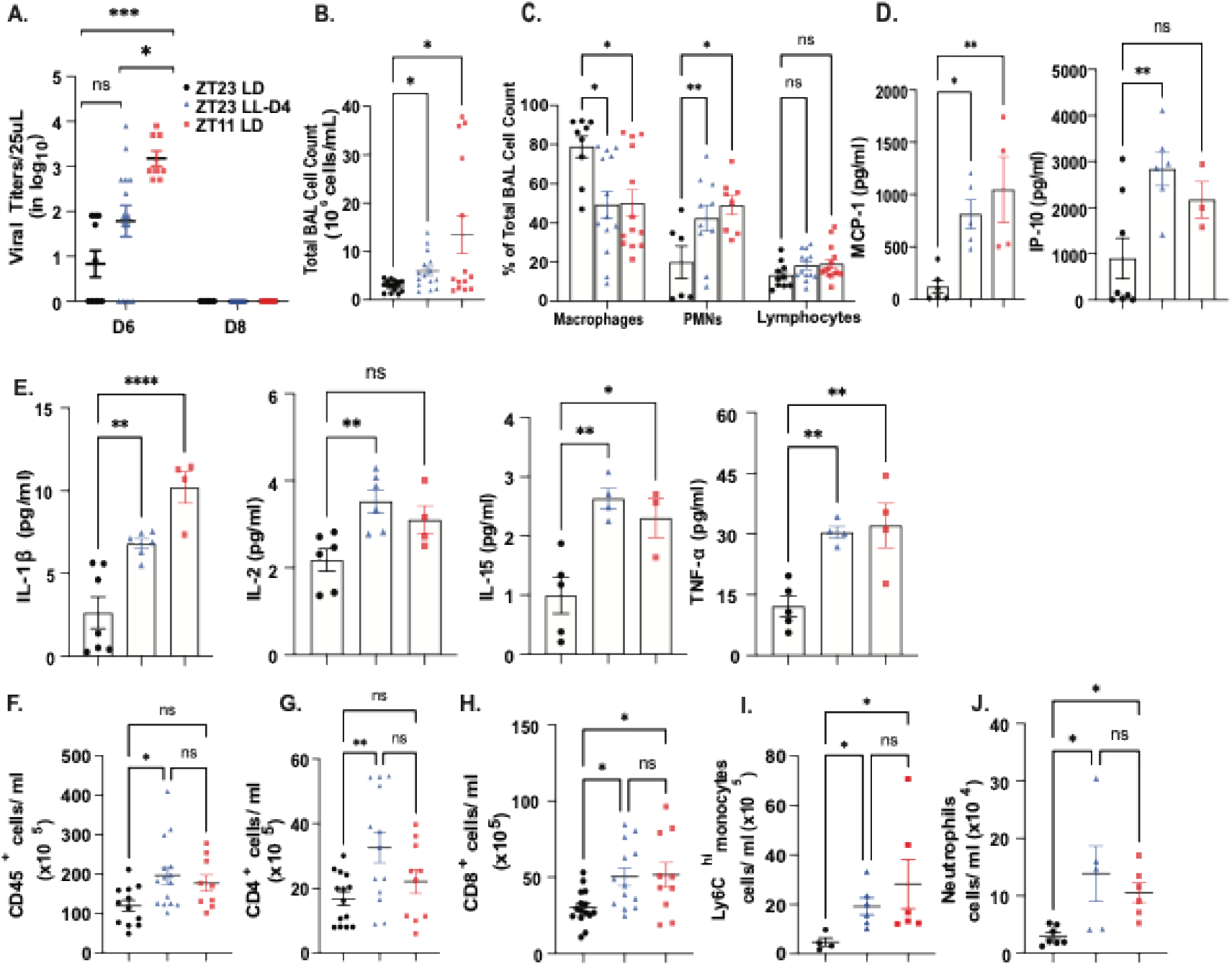
Exposure to constant light following IAV infection causes exaggerated inflammation independent of viral burden. (A) Quantification of viral titers (Day 6, n = 8-14 per group, Day 8, n= 6-10 per group ***p<0.0005; two-way ANOVA with main effects only) (B) Total Bronchoalveolar lavage (BAL) on 6 days p.i. (n = 13-15 per group, *p<0.05 one-way ANOVA; Kruskal Wallis test) (C) Differential of the total BAL cells (n = 6-13 per group, *p<0.05 ; **p<0.001 two-way ANOVA, mixed effects model) (D) Chemokine levels in BAL on day 6 p.i. (n = 3-8 per group, *p<0.05 ; **p<0.001 Ordinary One-way ANOVA) (E) Cytokine levels in BAL on day 6 p.i. (n = 3-7 per group, *p<0.05 one-way, **p<0.001, ****p<0.0001 Ordinary One-way ANOVA) (F) Absolute number of CD45^+^ cells (G) Absolute number of CD4^+^ T lymphocytes (H) Absolute number of CD8^+^ T lymphocytes (F-H) n = 9-15 per group; *p<0.05 one-way, **p<0.001 Ordinary One-way ANOVA (I) Absolute number of Ly6C^hi^ inflammatory monocytes and (J) Absolute number of Neutrophils. n = 4-7 per group *p<0.05 Ordinary One-way ANOVA; Kruskal Walli’s test. All data pooled from 3-5 independent experiments. ns= non-significant

### Constant light exposure worsens immune-mediated pathology in IAV infection

Given the viral titers were comparable in the ZT23(LL-D4) and ZT23(LD) groups despite significantly higher mortality in the former, next, we investigated if exposure to constant light worsened lung inflammation. Consistent with our previous work, the total Bronchoalveolar lavage (BAL) cell count was higher in the mice infected at ZT11(LD) than in those infected at ZT23 [Fig 2B]. Interestingly, the ZT23(LL-D4) group mice had higher total BAL cell counts than the ZT23(LD) group, comparable to the ZT11(LD) group [Fig 2B]. The higher BAL count in the ZT23 LL-D4 and ZT11 mice was accompanied with a significantly lower number of macrophages with higher total BAL cell counts had a significantly lower number of macrophages when compared to the ZT23 (LD) group [Fig 2C]. Next, to test if a short period of LD cycling-driven circadian disruption can result in a state of heightened inflammation in the lungs, we examined BAL from naïve mice -exposed to constant light for 4 days. We found no differences in the total BAL cell counts between the mice maintained in 12 hr LD cycling and those exposed to constant light [Fig 4H]. Considered together, this supports the idea that the effect of short periods of light disruption is exacerbated in an infected host. The ZT23(LL-D4) and ZT11(LD) groups had significantly more polymorphonuclear cells in the BAL than the ZT23(LD) group [Fig 2C]. Further, compared to the ZT23(LD) group, on day 6 p.i. the levels of chemokines, MCP1 and CXCL10, were significantly higher in the BAL of the ZT23(LL-D4) group, whereas ZT11(LD) had only higher levels of MCP-1 [Fig 2D]. The levels of cytokines IL1β and IL15 were also significantly higher in both the ZT23(LL-D4) and ZT11(LD) groups than the ZT23(LD) group, whereas the level of IL-2 was only significantly higher in the ZT23(LL-D4) [Fig 2E]. Interestingly, there was no significant difference in IFNγ levels across the three groups on day 6 p.i.[Fig 2E]. Overall, the chemokine/cytokine profile of the ZT23(LL-D4) group was aligned with the ZT11(LD) group. Based on these analyses, we hypothesized that the ZT23(LL-D4) group would have more pro-inflammatory immune populations than the ZT23(LD) group. Therefore, we harvested lungs on day 8 p.i. and compared the immune populations in the three groups. Consistent with our previous work and our BAL analyses, we found that the ZT23(LL-D4) mice had more leukocytes in the lungs than the ZT23(LD) mice [Fig 2F]. Since the ZT23(LL-D4) group had higher levels of MCP1 and CXCL10 --chemokines that promote monocyte and lymphocyte trafficking, respectively, into inflamed lungs, we enumerated these populations, Ly6C^hi^ inflammatory monocytes (CD45^+^ Ly6G^-^ CD11b^+^Ly6C^hi^) and T cells (CD45^+^CD4^+^ and CD45^+^CD8^+^), in the lungs. Both populations were higher in the ZT23(LL-D4) than the ZT23(LD). We were higher in the ZT23 (LL-D4) group compared to the ZT23-LD group [Fig 2G-I]. The absolute number of neutrophils (CD45^+^Ly6G^+^CD11b^+^) was also higher in the ZT23(LL-D4) group than in the ZT23(LD) group [Fig 2J]. Thus, the disruption of environmental lighting drives an exaggerated inflammatory milieu with higher levels of chemokines and inflammatory cell populations.

### Constant light exposure worsens lung injury from IAV

To validate the above results, we undertook histological assessment of lungs harvested from the three groups. Consistent with our previous results, using a validated scoring system^5,6^, the ZT23(LD) mice sustained significantly less injury than the ZT11 group [Fig 3A-I]. Lung pathology was markedly worse for the ZT23(LL-D4) than the ZT23(LD) group, with higher peri-bronchial infiltrates, peri-vascular infiltrates, inflammatory alveolar exudates, and epithelial necrosis [Fig 3A-I]. Further, we validated our results from the flowcytometric analyses on histology from day 8 p.i. by staining for lymphocytes(CD3^+^) and macrophages and monocytes (F4/80^+^). The lungs harvested from the ZT23(LL-D4) and ZT11(LD) groups had a significantly higher proportion of CD3^+^ and F4/80^+^ cells per high-power field (HPF) than that of the ZT23(LD) group [Fig 3C-I, & E-F respectively]. Next, we investigated if constant light exposure would affect lung regeneration, even beyond the acute immunopathology described above. After severe lung injury, alveolar type 2 (AT2) cells promote regeneration of the airspace or alveoli. We found that there were significantly fewer Alveolar type 2 (AT2) cells, denoted by SFTPC^+^ cells/HPF, in the lungs of the ZT23(LL-D4) and ZT11(LD) groups than in the ZT23(LD) group [Fig 3G-I]. This could affect how the lungs regenerate after the initial inflammatory response. Although the primary driver for the poorer outcomes in the ZT23(LL-D4) group, like that for the ZT11 group historically, is immune-mediated pathology, a loss of regenerative capacity would likely worsen recovery from IAV, leading to long-term morbidity.

**Fig 3.**
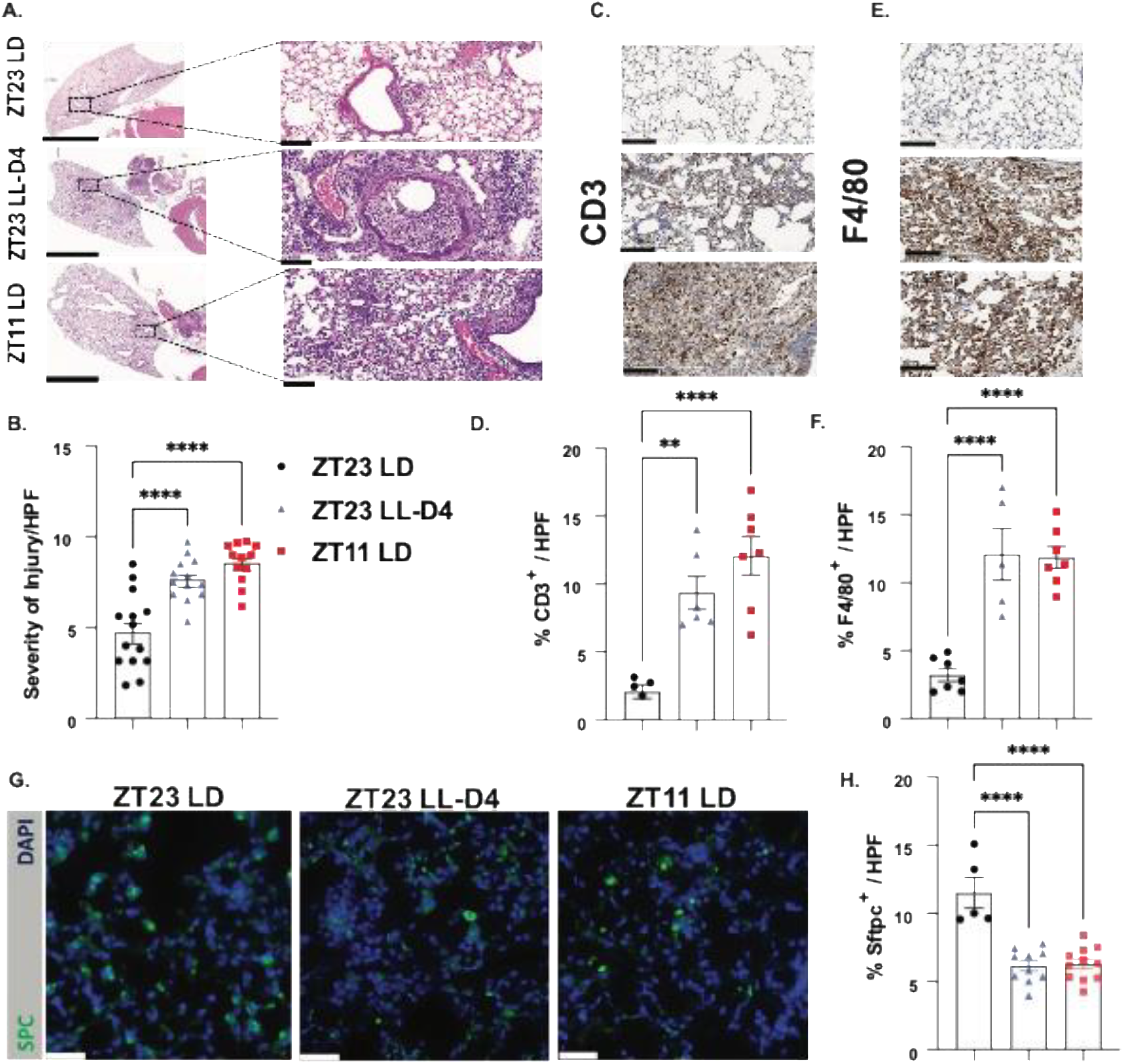
Immunopathology from IAV is worsened by environmental light cycling disruption (A) Representative micrographs of H&E-stained lung sections on day 8 p.i. Left insert: Scale bar = 5mm; Right insert: Magnification: 20X; Scale bar = 200 μm (B) Quantification of acute lung injury (n = 12-14 per group; ****p<0.0001 Ordinary One-way ANOVA) (C) Representative micrographs of CD3-stained lung sections on day 8 p.i. at 20X; Scale bar = 200 μm (D) Quantification of CD3^+^ cells per high field (HPF) (n = 5-7 per group; **p<0.001, ****p<0.0001 Ordinary One-way ANOVA) (E) Representative micrographs of F4/80-stained lung sections on day 8 p.i. at 20X; Scale bar = 200 μm (F) Quantification of F4/80^+^ cells per high field (HPF) (n = 5-7 per group; ****p<0.0001 Ordinary One-way ANOVA). (G) Representative images of pro-SPC-stained lung sections on day 8 p.i. at 40X (H) Quantification of SPC^+^ cells per high field (HPF) (n = 5-11 per group; ****p<0.0001 Ordinary One-way ANOVA). Scale bar = 10 μm. All data pooled from 5 independent experiments. ns= non-significant

### Light disruption is effective during a specific window of vulnerability following influenza A infection

Having established the need for synchronizing influence of entraining photic cues in maintaining the clock-driven protection from IAV in the early phase of influenza, we asked if there exists a window of vulnerability to light disruption following influenza infection. To test this, we infected mice at either ZT23 or ZT11 and maintained them in 12 hr LD cycling conditions until day 7 p.i. At this time, a subset of the ZT23 groups was moved to constant light, referred to as the ZT23(LL-D7), instead of day 4 in earlier experiments [Fig 4A].

**Fig 4.**
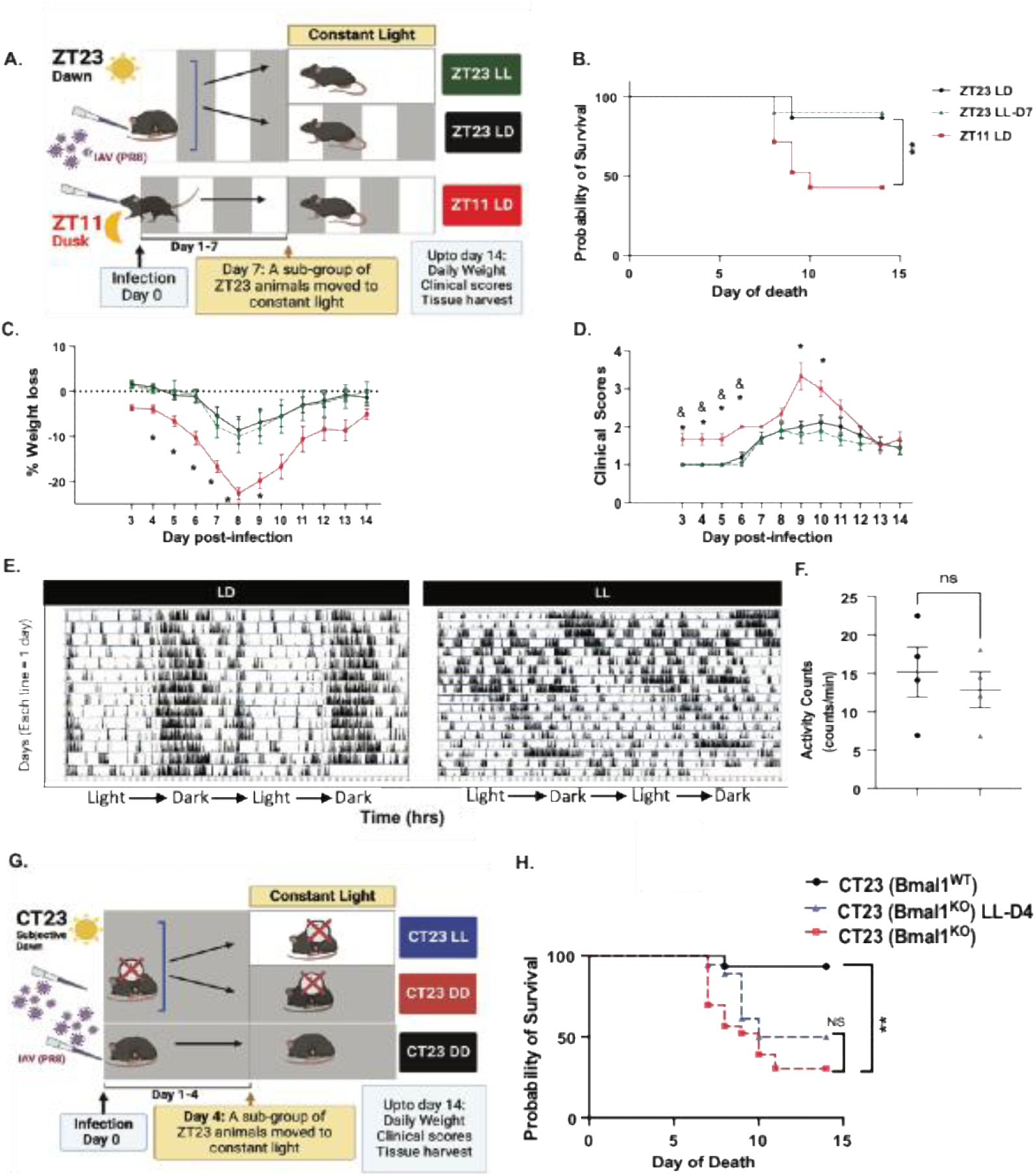
A specific window of vulnerability to light disruption following influenza A infection exists (A) Experimental model (B) Survival (n = 10–21 per group, **p<0.001 log-rank test) (C) Weight loss trajectory (n=10-21 per group, *p<0.05 ANOVA for repeated measures) following IAV infection (D) Average clinical score. (n=10-21 per group, *p<0.05 ANOVA for repeated measures). * = ZT23 vs ZT11; & = ZT23 LL-D7 vs ZT11. (E) Representative double-plotted actogram of naïve mice maintained under 12-hour light-dark cycling followed by constant light conditions; LL (n=4) (F) Characteristics of circadian rhythm, Amplitude *p<0.05; unpaired t-test; Welch’s corrrection (G) Average activity on actograms (H) Left: Total BAL cell counts from naïve mice maintained either in 12 hr light dark conditions or exposed to constant light for 5 days. Right: Differential of total BAL cell counts. (n = 4-8 per group, Ordinary One-way ANOVA; mixed effects model) (I) Experimental model (J) Survival (n = 10–18 per group, **p<0.001 log-rank test). All data pooled from three independent experiments. ns= non-significant

The ZT23(LD) group had significantly lower mortality than mice infected at ZT11(LD). However, unlike the results from constant light exposure on day 4 p.i., the time-of-day difference in outcomes was not lost when animals were moved to constant light on day 7 p.i. [Fig 4B: 86.66% survival in ZT23(LL-D7) vs 90% in ZT23(LD) versus 42.8% in ZT11(LD)]. Further, the weight loss trajectory and morbidity of the ZT23(LL-D7) group was comparable to that of the ZT23(LD) group [Fig 4C-I], suggesting that there is a window of vulnerability to LD disruption following influenza infection.

### The central clock mediates the abrogation of time-of-day specific protection

To investigate if the central clock mediates the loss of time-of-day-specific protection in influenza infection in mice exposed to constant light, we analyzed the locomotor activity as a read-out for the former. Since singly housing mice (necessary for locomotor activity monitoring) after IAV infection could worsen outcomes, we used naïve mice in this experiment. Mice were exposed to 4 weeks of LD cycling and then moved to constant light conditions for 3 weeks. We found that the mice in LD cycling showed clear demarcation of active and rest phases coinciding with dark and light phases of the 24-hour cycle [Fig 4E]. The mice exposed to constant light display having erratic onsets of activity, eventually becoming completely arrhythmic after being exposed to constant light for three weeks [Fig 4E]. Mice kept in continuous light although arrhythmic had comparable activity levels, supporting the idea that constant light led to loss of rhythmicity but did not impair overall activity level [Fig 4F-I]. Prolonged constant light exposure is also associated with increased stress response^7,8^. However, given our context-specific and short duration of LL exposure, we hypothesized that the effect of LL in our influenza model mediates its effect mainly through circadian disruption and not additional stress independent of circadian disruption.

To test this, we used the *iBmal1*^*-/-*^ mice (*Bmal1*^*fl/fl*^ *ERT2Cre*^*+*^), where the clock has been genetically disrupted globally; these mice are arrhythmic in DD conditions^9^. Previously, we have showed that these *iBmal1*^*-/-*^ mice do not have clock-driven time-of-day specific protection against IAV^5^. Initially, we tested the effect of LD cycling on WT and *iBmal1*^*-/-*^ mice to keep it consistent with our experimental design as outlined in Fig 1. However, we found that *Bmal1*^*-/-*^ mice under LD conditions showed a reversal of the time of day-specific protection compared with the WT mice, with *iBmal1*^*-/-*^ mice infected at ZT11 having better outcomes than *iBmal1*^*-/-*^ mice infected at ZT23 (Supplementary Fig 1). This would suggest that in the absence of the molecular clock LD cycling primes processes allowing a reversal, but not a abrogation of the time-of-day protection. Thus, based on our previously published work, we tested the *Bmal1*^*-/-*^ mice in constant darkness (DD) conditions^5^. Generally speaking, CT time refers to the same times the as ZT but under DD conditions. Therefore, CT23 represents subjective dawn. We infected *Bmal1*^*-/-*^ mice and their cre^neg^ littermates maintained in DD conditions. On day 4, p.i., a subset of *Bmal1*^*-/-*^ mice were moved from constant darkness to constant light conditions [Fig 4I]. We found that consistent with our previous results, *Bmal1*^*-/-*^ mice infected at CT23 experienced significantly higher mortality than wild-type littermates [p<0.001, Mantel-Cox log-rank test, Fig 4J]. However, while *Bmal1*^*-/-*^ mice exposed to constant light still had higher mortality than the WT littermates, there was no difference in the outcomes of *Bmal1*^*-/-*^ mice in the two lighting conditions [Survival of 31% in *iBmal1*^*-/-*^CT23 and 50% in *iBmal1*^*-/-*^ CT23(LL-D4)], suggesting that constant light did not worsen the outcomes over that attributed to loss of the clock.

### Global immune activation observed in the dawn-infected group when exposed to constant light post-infection

To determine what pathways resulted in worse outcomes in the ZT23(LL-D4) group, we performed transcriptional profiling on lungs harvested on day 8 p.i from both ZT23(LD) and ZT23(LL-D4) (n = 5/ group; all females) [Fig 5A]. Environmental light disruption following influenza infection resulted in disparate transcriptional phenotypes [Fig 5B]. 1125 genes were differentially expressed between the two groups, with 801 genes upregulated and 324 genes downregulated in ZT23(LL-D4) relative to the ZT23(LD) group. Several genes associated with the migration of leukocytes (*Ccr5, Ccl4, Cxcl10*) into the lung and exaggerated immune activation (*Il18rap, Gzma, Ly6c2*), were upregulated in the ZT23(LL-D4) group compared with the ZT23(LD) group [Fig 5C]. Further pathway analysis revealed that while those involved in innate activation (“Granulocyte diapedesis”, “IL15 signaling”, “Cytokine storm”) were upregulated in the ZT23(LL-D4) group relative to the ZT23(LD) group, the most highly significant pathways involved innate to adaptive communication (“Communication between innate and adaptive cells”) and activation of the adaptive immune system (“B cell activation as in “Systemic lupus”, “Altered T and B cells signaling”, “NFAT signaling”) [Fig 5D]. Overall, these pathways supported our findings of excessive immune cells, especially lymphocytes and monocytes, in the lungs of the ZT23(LL-D4) group compared with the ZT23(LD) group. Thus, even a short duration of light-based circadian disruption causes profound alteration of the immune response.

**Fig 5.**
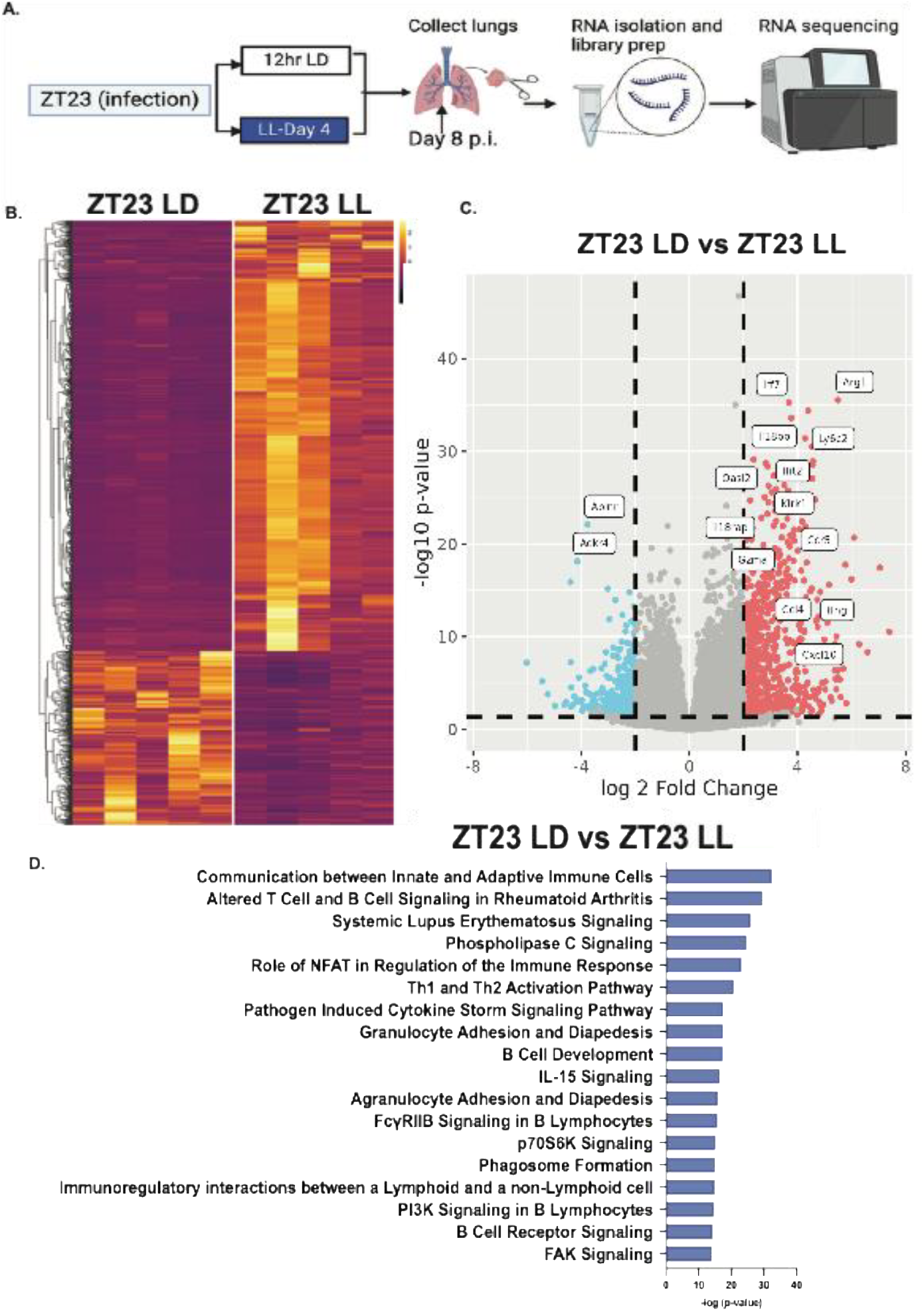
Transcriptomic analyses show global immune activation in ZT23 LL and ZT11 LD groups. (A) Schematic of bulk RNA-seq processing on Day 8 p.i. lungs. (B) Heatmaps of all differentially regulated genes. (C) Volcano plots showing upregulated and downregulated genes. In each volcano plot, the horizontal dotted line represents a Padj = 0.05 and the vertical dotted line represents a log (fold change) >2 or <-2. (D) Plot of log-adjusted fold change for ZT23 LD vs ZT23 LL showing directionality of the most differentially expressed genes. n = 5 per group, all females. All data pooled data from three independent experiments.

### Transcriptomic parallels between the effect of light disruption and time-of-day effects

While we have shown that the ZT11(LD) and ZT23(LL-D4) groups both show excessive immune infiltration and subsequent tissue destruction[Fig 2 and 3], we wanted to test if the mechanism(s) underlying the poorer outcomes in both these groups were indeed similar. For this, we expanded our transcriptomic study to include three groups-ZT23(LD), ZT11(LD), and ZT23(LL-D4) [Fig 6A]. There were 1304 differentially expressed (DE) genes in comparing the ZT23(LD) with the ZT11(LD) group. Of these, 856 genes were upregulated and 448 genes were downregulated in the ZT11(LD) group relative to the ZT23(LD). Of the 1304 DE genes, 739 genes were shared with the comparison between ZT23(LD) vs ZT23(LL-D4) (Fig 6A). Of these 739 shared DE genes, 587 genes were upregulated, while 151 genes were downregulated in both ZT23(LL-D4) and ZT11(LD) relative to ZT23(LD). Only one low-expression gene (*Oxtr*) was downregulated in the ZT23(LL-D4) and upregulated in ZT11(LD). Thus, not only was there significant overlap in the DE genes, but their directionality was also conserved across the two comparisons – that of ZT23(LL-D4) versus ZT23(LD) and ZT11(LD) versus ZT23(LD), respectively. Next, we compared the ZT23(LL-D4) with the ZT11(LD) group to further confirm the mechanistic overlap. We found only 13 DE genes with ≥ a 2-fold change in expression between these two groups; of these, many were circadian genes like *Bmal1(Arntl), Dbp, and Npas2*, whose expression oscillates across the day and is expected to be disrupted by constant light. In further analyses, as expected, various innate immune pathways, namely, “cell adhesion and diapedesis” and “the role of hypercytokinemia/ hypercytokinemia in the pathogenesis of influenza” are enriched and conserved in both comparisons [Fig 6C]. Pathways involved in adaptive immune activation were also likewise enriched in the ZT23(LL-D4) and ZT11(LD) groups relative to ZT23(LD) [Fig 6C]. Genes involved in leukocyte migration (*Ccl4, Ccr5*) and activation of the immune system (*Ifng, Klrk1,Gzma, Il18rap*) were all higher in the ZT23(LL-D4) and ZT11(LD) groups compared to ZT23(LD). We confirmed changes in expression in some of the key genes by qPCR [Fig 6D-I]. Together, these results suggest that LD-cycling perturbation following influenza impairs lymphocyte and monocyte diapedesis and results in immunopathology similar to that seen the clock-regulated less favorable time of day. In other words, processes affected by light disruption usually are regulated by the clock to confer time-of-day-specific protection.

**Fig 6.**
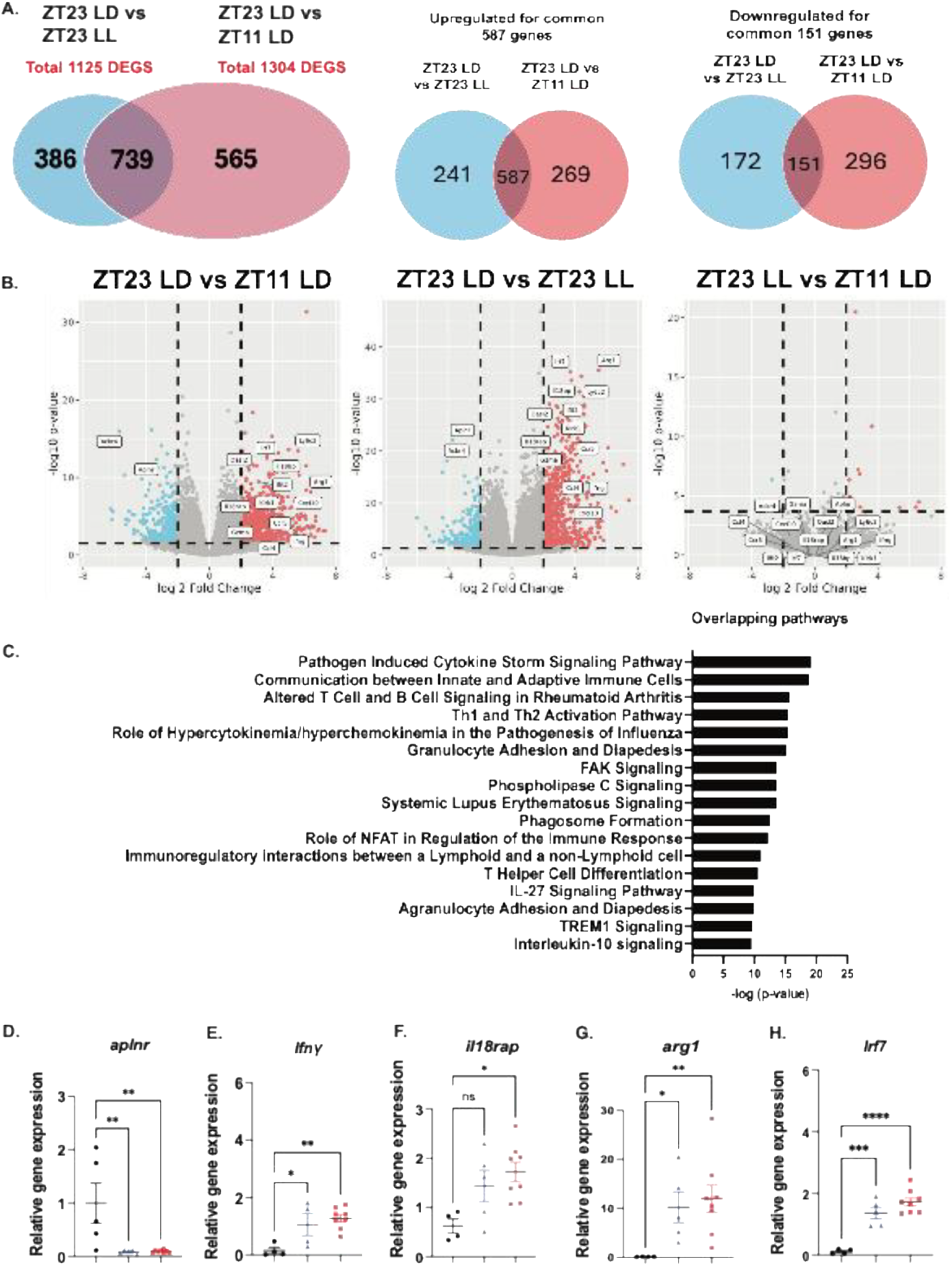
Transcriptomic analyses show global immune activation in ZT23 LL and ZT11 LD groups. (A) Venn diagram (sizes not to scale) depicting the number of differentially expressed genes. (B) Volcano plots showing upregulated and downregulated genes. In each volcano plot, the horizontal dotted line represents a Padj = 0.05 and the vertical dotted line represents a log (fold change) >2 or <-2. (C) Plot of pathway enrichment p-values for overlapping genes differentially expressed both between ZT23 LD vs ZT23 LL and between ZT23 LD vs ZT11 LD. (D-H) Relative gene expression of selected genes as a confirmation for bulk RNA seq. n = 4-8 per group, *p<0.05, **p<0.001, ***p<0.0005, ****p<0.0001; Ordinary One-way ANOVA. All data pooled from three independent experiments; ns = non-significant.

### Cycling of food availability restores the time-of-day specific protection from influenza A infection in the face of environmental light disruption

While light is the primary external cue that entrains and synchronizes the central and peripheral clocks, other external cues like exercise and meal timing may significantly affect peripheral clocks^10^. To investigate if the poor outcomes caused by environmental light perturbation could be rescued by using a different entrainment cue, we considered a role for food cycling (FC) in the presence of environmental light perturbation. We followed the experimental design as in Fig 1A, where mice were infected at either ZT23 or ZT11, and a subset of mice infected at ZT23 were moved to constant light on day 4 post-infection. Within the ZT23(LL-D4) groups, a subset of mice was subjected to food cycling, wherein food was provided from ZT12 to ZT0 each day [ZT23(LL-D4:FC)] while the other had access to food ad libitum [ZT23(LL-D4:AL]; all groups had unrestricted access to water [Fig 7A]. Consistent with our previous results, the ZT23(LL-D4) with food ad libitum group had higher mortality than the ZT23(LD) group. As in previous experiments, the ZT23(LD) group had significantly better survival than the ZT11(LD) group [Fig 7B: 87% survival in ZT23(LD) versus 37.5% survival in ZT11(LD); p <0.001 by Mantel-Cox Log-rank test]. Likewise, the exposure to constant light significantly reduced the survival in the ZT23(LL-D4) [Fig 7B: 51% survival in ZT23(LL-D4) versus 86% survival in ZT23(LD) group. P<0.05 by mantel-Cox Long-rank test]. Interestingly, the mice from the ZT23(LL-D4:FC) group showed rescue of the survival advantage associated with infection at ZT23 that had been lost in the ZT23(LL-D4: AL) group [Fig 7B: Survival of 81% in ZT23(LL-D4:FC) vs. 51% in ZT23(LL-D4); p<0.05, Mantel-Cox Log-rank test]. The mice in the food-cycling group also lost less weight than the ZT23(LL-D4:AL) group. [Fig 7C]. And the ZT23(LL-D4:FC) groups also exhibited lower clinical scores, suggesting less morbidity than the ZT23(LL-D4:AL) group [Fig 7D]. These findings lead to the conclusion that while LD cycling and meal timing are important zeitgebers affecting the host’s circadian rhythms, the latter is hierarchically a more potent entraining factor for optimal host response to influenza infection.

**Fig 7.**
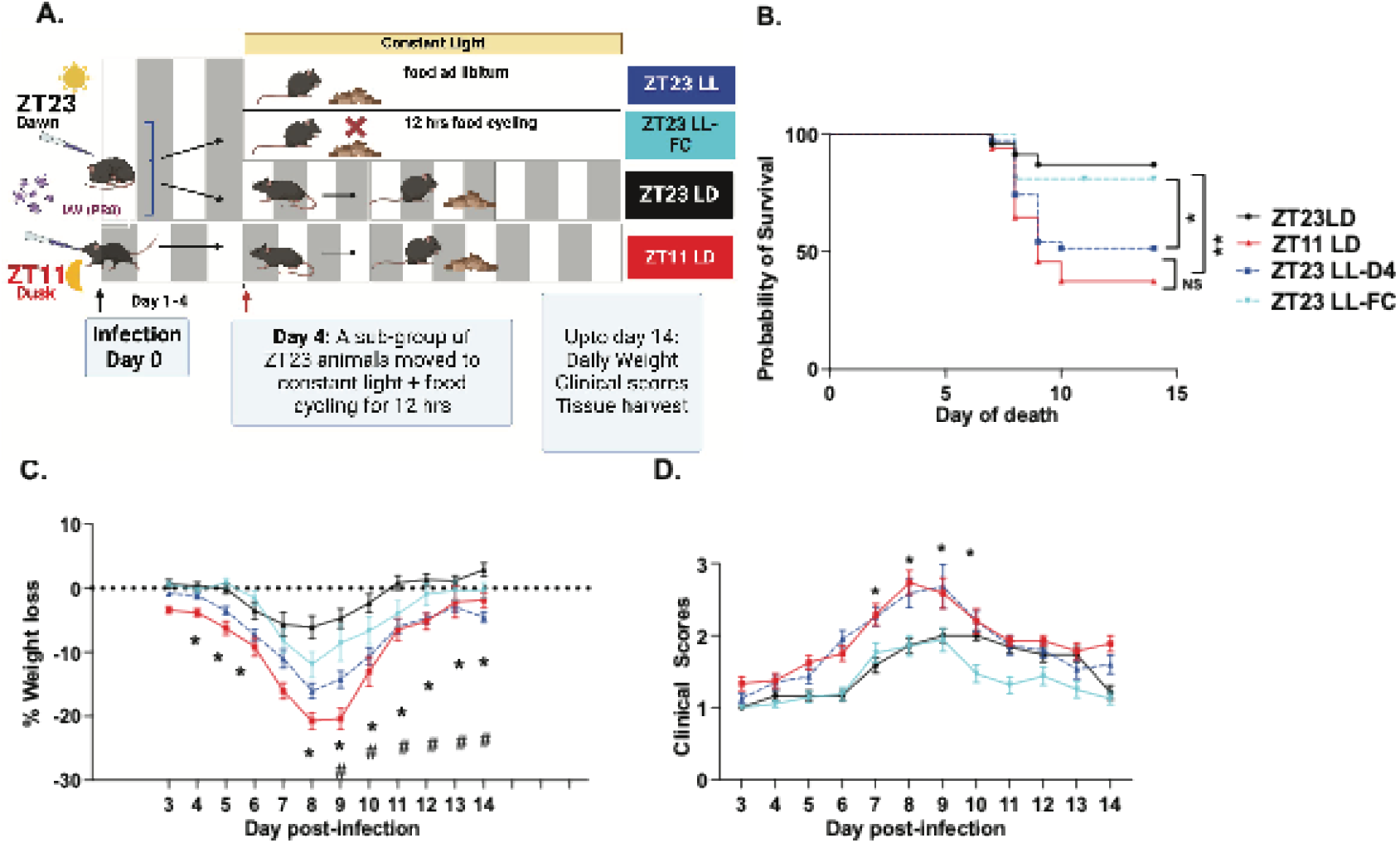
Food cycling rescues the loss of time-of-day protection from influenza A following environmental light disruption. (A) Experimental model (B) Survival (n = 21-50) per group, **p<0.01 log-rank test from three independent experiments, *p<0.05 log-rank test from three independent experiments) (C) Weight loss trajectory (n= 21-50 per group, *p<0.05; ANOVA for repeated measures) following IAV (D) Average clinical score. (n=21-50 per group, *p<0.05 ANOVA for repeated measures) following IAV infection. All data pooled from three independent experiments * ZT23 LD vs ZT11 LD; # ZT23 LD vs ZT23 LL-D4; NS = Non-Significant

## Discussion

While we and others have demonstrated time-of-day specific protection conferred by the circadian clock^5,11,12^, our study uncovers how common zeitgebers interact dynamically after the initial infection to shape the host’s response to the pathogen. Circadian disruption is known to worsen outcomes for many health conditions, including infection, most often, these are genetic or environmental disruptions that predate the infection^12-15^. When time-of-day studies are undertaken for infections or other discrete stimuli, the sentinel event is deemed to occur on the day of the infection. It is unknown whether circadian mechanism(s) that directs host protection is significant, in the aftermath of an infection.

We have previously showed that mice infected at ZT23 had a 3-fold higher mortality rate than mice infected at ZT11^5^. Since the peak mortality from influenza infection is between days 8-10 in mice, questions have remained on whether it is the early signaling following infection or an effective but continuing cascade of well-regulated immune response to IAV that drives the time-of-day specific protection. While the effect of longstanding disruption of LD cycling as simulated by chronic jetlag on health outcomes^16^ is well known, the impact of short-term disruption of LD cycling after acute infection is poorly understood. Our study shows that this time-of-day-specific protection in the ZT23 group is lost when mice are subjected to circadian disruption via constant light exposure. The effect of circadian disruption, as seen with constant light exposure, has been studied in other contexts^17,18^. However, the role of a short duration of light exposure on acute lung injury has not been investigated before. Some studies, although not investigating the lungs, used > 4 weeks of constant light exposure to define outcomes^19,20^ or higher-intensity light cycling at baseline^21^. We found that even a short duration of constant light exposure in an infected animal worsened lung inflammation.

Our locomotor activity analyses documenting the disruptive effect of constant light on circadian rhythms were done in the naïve host and with a significantly longer duration of constant light exposure [Fig. 4E]. We found it technically challenging to house mice singly^22,23^ for sustained periods (that would be necessary for doing locomotor activity analyses), especially given the trajectory of the acute illness of influenza infection. Although this protocol clearly demonstrated loss of rhythmicity, we verified the role of the clock in the constant light paradigm by conducting light disruption in a clock-mutant background. We showed that in mice with genetically disrupted clocks constant light exposure did not worsen outcomes following IAV. Our work on the impact of LD cycling on the *Bmal1*^*fl/fl*^ *ERT2Cre*^*+*^ mice also highlights the complexity of the entrainment against a background of lack of endogenous clock in the clock-disrupted mice.

Importantly, we show that food cycling-based entrainment was able to restore the clock-driven advantage lost due to disruption of photic cues in influenza infection. This would support the idea that metabolic cues are more critical for the entrainment of peripheral clocks than photic cues when facing an energy-demanding function like fighting an infection. The responsiveness of tissue-specific clock to food cycling has been demonstrated for the liver clock^18,24-26^ and is supported by reports from the lung^27,28^; however, these studies involved longer entrainment and were not performed in infected hosts. Finally, based on the short duration of constant light and food cycling that changes the trajectory in the infected mice infected at ZT23, we speculate that maintaining “entraining cues” for the endogenous circadian clock is especially critical in an infected host. Given the acute immune and metabolic stress from the infection, such a host may be especially vulnerable to circadian perturbation.

Our model is of particular interest for its translational relevance. Chronic circadian disruption, as seen in shift workers, has been associated with increased risk for cancers, hypertension, diabetes mellitus, and metabolic syndrome^4^. However, circadian disruption is widespread in hospitals and intensive care units^29-31^. LD cycling is frequently disrupted in these units, with dim and limited-spectrum light exposure during the day and erratic but more light exposure at night^32^; while such disruption has been linked to delirium or other neuropsychiatric symptoms, to our knowledge we are the first to show deleterious effects at the pulmonary level. Given the challenges of simulating erratic LD cycling, we used constant light for these proof-of-concept studies and uncovered pathogenic consequences of disrupted rhythms a few days after infection. Our work furthers our understanding of the host circadian rhythms needed to maintain clock-gated protection in the aftermath of respiratory illnesses and suggest non-invasive interventions to restore cycling of external cues to boost circadian health of the host. Considering these entraining factors in the clinical settings offers a new avenue for improving outcomes following severe viral infections.

## Acknowledgments

We are grateful to Shivani Rawat and Paine Fleisher for their help with animal husbandry. This work was supported by NHLBI R01HL155934-01A1(SS) and NHLBI-R01HL147472 (SS). A Shetty was supported by CURF and AC by the PURM grant, University of Pennsylvania. A Sehgal is an HHMI investigator. We are also grateful to Christoph Thaiss for helpful discussions.

## Author Contributions

SS conceived the project; SS, OP and ASehgal designed experiments; OP, KF, LA, MT, A Shetty, and SS performed experiments and collected data. SS, OP, AC, A Setty, LA, and TB analyzed data. GG was involved in the analyses and interpretation of the transcriptomic data. OP, SS, and TB wrote the original draft, and A Sehgal and GG helped with revisions. SS supervised all research activities.

## Data Availability

The RNA-Sc-seq data is available at the NCBI GEO accession number GSE288858. All other data is available as source data.

## Methods

### Mice, virus, infection, constant light exposure, and food cycling

Specific pathogen-free 8-12 week-old C57bl/6J mice (both males and females) were placed in circadian cabinets on reverse light-dark cycles as described previously (citation). This allowed us to simultaneously infect mice or harvest tissues across all experimental groups. After 4-6 weeks of acclimatization in these cabinets, mice were anesthetized lightly with isoflurane prior to being infected intranasally (i.n.) with mouse-adapted influenza virus (PR8). Given the use of circadian cabinets, mice were simultaneously infected at either ZT23 or ZT11 and maintained in 12 hr light-dark cycle. The dose chosen was likely to cause >50% mortality when mice were infected at ZT11. Where relevant, on days four or seven, p.i. a subset of ZT23-LD mice were moved to constant light until the end of the study. Animals were weighed and scored based on a previously validated scoring criterion each day. For a subset of the ZT23 LL mice, the food was cycled to a 12-hour period, coinciding with their active and rest phases, which had them remain in 12-hour LD conditions. Food trays were manually removed for the food cycling experiments, and the overall food consumed was not monitored in any group to avoid housing mice singly after IAV infection. The food trays in non-food cycled study groups were handled but not removed to control for any stress associated with this procedure. Approval from the Children’s Hospital of Philadelphia Institutional Animal Care and Use Committee was obtained for all animal studies, and the stipulations were met according to the Guide for the Care and Use of Laboratory Animals. Male and female mice were used in approximately equal proportions for individual experiments unless specified. Only female mice were used for the RNA-seq experiment to minimize any variability due to sex-specific effects.

### Genetic mouse mutants

Inducible Bmal1 knockout mice were generated using 2-month-old *Bmal1*^*fl/fl*^ *ERT2Cre*^*+*^ mice treated with 5mg (in 50 ul) tamoxifen via oral gavage daily for 5 consecutive days^5^. *Bmal1*^*fl/fl*^ *Cre*^*neg*^ littermates treated with tamoxifen were used as controls. They were exposed to constant darkness for 1-2 days before infection with IAV at CT23. Both males and females were used in equal proportions in the mentioned experiment.

### Viral titration

Lungs were harvested and homogenized in 0.1% PBS-gelatin. Influenza virus was detected by using MDCK cells (gift from Scott Hensley’s group: originally purchased from ATCC, cat no. PTA-6500). MDCK cells were overlaid with 1:10 dilutions of the homogenized lungs and incubated at 37^°^C; 5% CO_2_ for 1 hr to allow the virus to infect the MDCK cells. Thereafter, 175 μl of tissue culture medium supplemented with 2 μg/ml of TPCK-treated trypsin was added, and cells were incubated for another 72 hrs at 37°C; 5% CO_2_ before viral titration. After 72 hrs, hemagglutination of turkey red blood cells (RBCs) was tested by collecting 50 μl of the medium from the infected MDCK plate to detect and quantify the virus. The hemagglutination of RBCs confirmed the presence of the virus particles^5^.

### Lung Histology, Immunofluorescence, and Bronchoalveolar lavage (BAL) cytology

A 20G flexible catheter (Surflo, Terumo, Philippines) was used to cannulate the tracheas. Using 600 μl of PBS, the lungs were lavaged gently in four passes. The supernatant from the first pass was collected and flash-frozen for ELISA. The cells from all four passes were pooled and counted using a Nexcelcom cell counter. Cytospins of BAL cells were stained with Hemacolor g(Sigma-Aldrich, Cat # 65044-93)). The lungs were perfused with 10 ml PBS and heparin, followed by inflation with 10% buffered formalin and then fixing in 10% buffered formalin for 24-48 hrs. The pathology core at Children’s Hospital of Philadelphia paraffin-embedded and stained the lungs for H&E stain. An Aperio CS-O slide scanner (Leica Biosystems, Chicago, IL) was used to digitally scan the stained slides at 20X by the pathology core. Aperio ImageScope v12.4 was used to take representative images of the scanned slides. Using a previously validated scoring method (Sengupta et al, 2019), scoring was performed with scorers remaining blinded to the group assignment. Briefly, five lung fields selected at random at 20X magnification were scored based on a scale of 0-3 for the following: 1) peri-bronchial infiltrates, 2) peri-vascular infiltrates, 3) alveolar exudates and 4) epithelial damage of medium-sized airways.

Paraffin-embedded lung sections were stained for macrophages and monocytes (F4/80) and lymphocyte (CD3) populations using immunohistochemistry (IHC) by the CHOP pathology core. The primary antibody used for myeloid staining was F4/80 (D2S9R) XP Rabbit mAB (Cell Signaling; Catalog # 70076) and Recombinant anti-CD3 epsilon antibody (EPR20752) (abcam; Catalog # ab215212) was used for staining the lymphoid population. All stained slides were digitally scanned at 20X using an Aperio CS-O slide scanner (Leica Biosystems, Chicago, IL). The lungs were annotated and analyzed using the Nuclear v9 macros from Aperio Imagescope software for F4/80 and CD3 analysis. Quantitation was performed using the Aperio Imagescope software. Data represent percent positive nuclei (for F4/80 and CD3^+^) per high power field (HPF). For immunofluorescence staining, the paraffin-embedded lung sections were treated in an oven at 60°C for an hour. The sections were incubated in xylene for 20 mins twice, then rehydrated in ethanol washes (100% twice, 90%, 80% & 70% ethanol) for 10 minutes each. Following antigen retrieval, slides were incubated in blocking buffer (5% donkey serum in PBST) for an hour at room temperature and then stained with primary antibody SFTPC (1:100, anti-pro surfactant protein, rabbit, AB3786 Millipore) at 4°C overnight. Post overnight incubation, slides were washed with PBST and incubated in secondary antibody diluted in PBST for an hour in room temperature. The secondary antibody used was at 1:250: Donkey anti-rabbit IgG (H+L) Alexa Flour 488 (Cat# A21206). After washing the slides with PBST, the slides were incubated with DAPI (0.2ug/ml) (Thermo Fisher; Cat#1738176) for 10 minutes. VectaMount AQ (Vector Laboratories Inc; Cat # H-5501) mounted the slides.

The histological and cytological scoring of all the samples was performed in a blinded fashion. The identity was unmasked once all the data were recorded, and final analyses were performed according to the study group. To quantify SFTPC^+^ cells post-influenza, images were captured at 20X on a Leica DM6 fluorescent microscope, and then cells were counted from >4 random sections per sample. The “Cell Counter” plug-in on ImageJ was used to count SFTPC^+^ cells. In total, >1000 cells were counted from n= 3-5 per group.

### ELISA

BAL from the first pass as described above on day 6 p.i. or lung homogenate in 0.1% gelatin on day 8 p.i. was sent for ELISA analysis to the Human Immunology Core at University of Pennsylvania for 9-plex cytokine array analysis. MCP-1, IP-10, IL-2, IL-1β, IL-15, TNF-α were measured using a custom ELISA kit (Milliplex Mouse Cytokine/Chemokine Magnetic Bead Panel; Cat # MCYTOMAG-70K-09C; Millipore SIGMA). Chemokines and cytokines were measured in 25 μl of BAL per sample in duplicates and the protocol for 9-plex Luminex Performance Assay was performed according to manufacturer’s protocol (Miilipore SIGMA). FLEXMAP 3D (BIO-RAD) was used to acquire data. Luminex x Potent 4.2 and Bio-Plex Manager 6.1 software (Bio-Rad) was used to analyze data.

### Locomotor Activity

Wireless infrared motion sensors (Actimetrics) were used to measure locomotor activity of singly housed mice before IAV infection. Once the uninfected mice were acclimatized to the circadian cabinets, their activity was measured in 12hr light-dark cycles for 4 weeks. After 4 weeks, the mice were exposed to constant light for another 3 weeks, and actogram data was collected. Parameters of the animals’ rhythms like amplitude, period, phase, average activity count and percentage of variance were analyzed using Clocklab software.

### RNA extraction and Bulk RNA sequencing

TRIzol (Life Technologies) was used to extract and isolate RNA from the inferior lobe of the mouse lung. The RNeasy Mini Elute Clean Up Kit (Qiagen) was used to further purify the RNA as per the manufacturer’s protocol. The NanoDrop ND-1000 spectrophotometer (NanoDrop Technologies Inc) was used to assess the quality and quantity of the extracted RNA. Only samples with RINS>7 after QC were sent for sequencing to Azenta Life Sciences. Sequencing was done using an Illumina NovaSeq X with Poly A selection to generate 2x 150 strand specific paired end reads. 20 million paired-end reads were obtained per sample. A mouse reference genome on an in-house resampling-based normalization and quantification pipeline aligned the samples. They were further compared to existing gene annotations (ENSEMBL) and novel loci and isoforms were identified. A false discovery rate-based control for multiple testing was used to identify differentially expressed genes. Finally, to assess fully the effects on key pathways and mediators, Ingenuity and GSEA were used.

### Flow cytometry

As described previously, the lungs were digested using DNAse II (Roche) and Liberase (II) at 37°C for 30 mins after perfusing the lungs with PBS through the right ventricle. The dissociated tissue was then passed through a 70μm cell strainer, followed by centrifugation and RBC lysis using ACK lysis buffer. After washing and resuspending the cells in PBS, 3 × 10^6^ cells were blocked with anti-CD16/32 antibody and first stained with Live/Dead fixability dyes for 30 mins. Following washing, the cells were stained with desired antibodies on ice for 30 mins. Cells were fixed in 2% paraformaldehyde. Flowcytometric data was obtained using LSR Fortessa cytometer and analyzed using FlowJo software (Tree Star, Inc). All cells were pre-gated on size as singlet, live cells. All subsequent gating was done on CD45^+^ cells only.

### Statistics

GraphPad (Prism V9 and V10) were used for all statistical analyses. A one-way or two-way ANOVA was performed in experiments with more than two groups and normally distributed data, Sidak’s correction was used for multiple comparisons. When data was not normally distributed, we used Mann-Whitney or Kruskal-Wallis tests. Most data is represented as mean ±SEM.

### Statement on rigor and reproducibility

Mice were either purchased from Jackson Labs or bred in-house. Reported findings are summarized results from 3-6 independent experiments.

## Notes

**Conflict of interest statement:** The authors have declared that no conflict of interest exists.

### Competing Interest Statement

The authors have declared no competing interest.

