## Supplemental Table 2 for "Effect of external cues on clock-driven protection from Influenza A infection"

### Slide 1
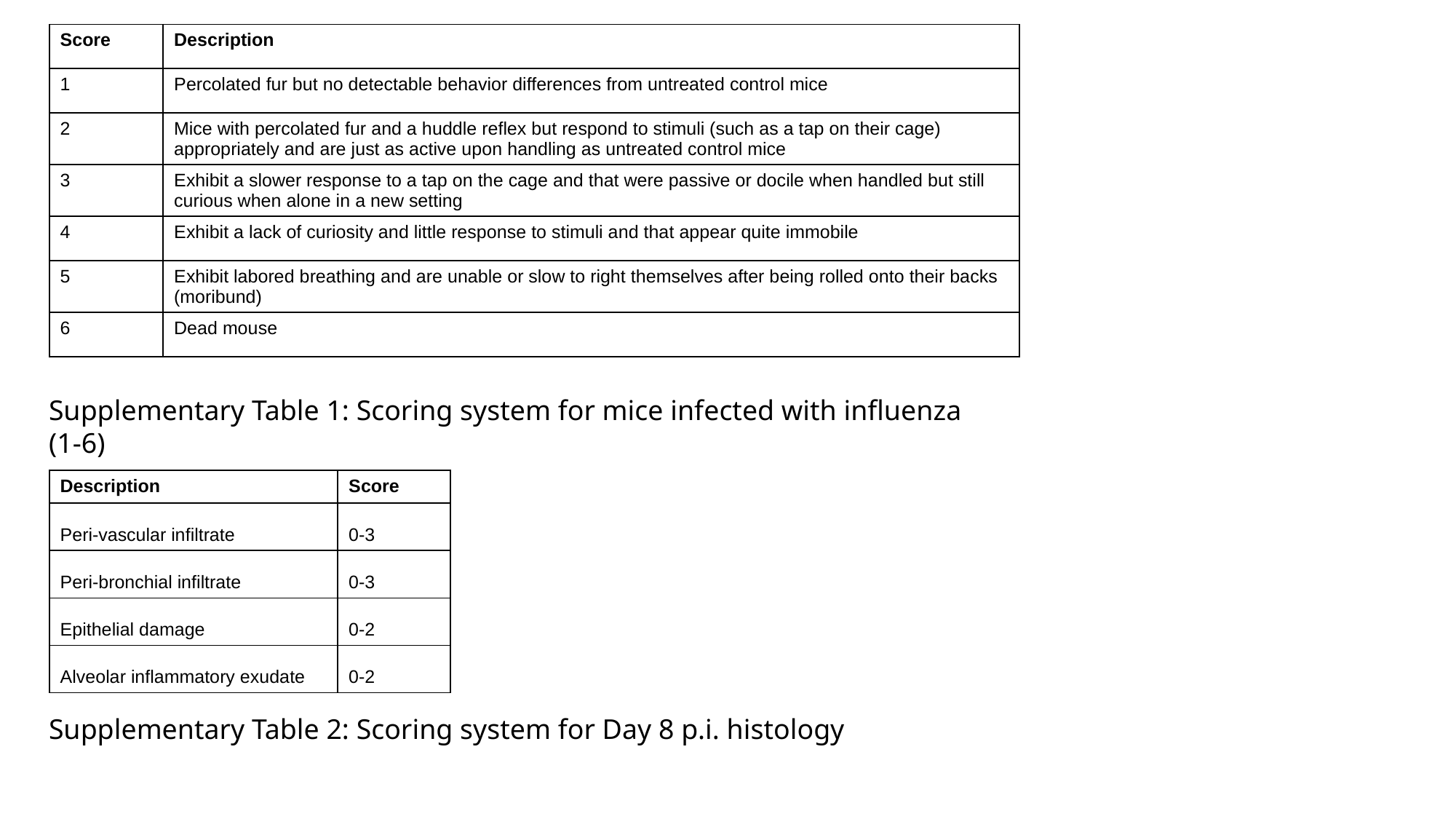

| Score | Description |
| --- | --- |
| 1 | Percolated fur but no detectable behavior differences from untreated control mice |
| 2 | Mice with percolated fur and a huddle reflex but respond to stimuli (such as a tap on their cage) appropriately and are just as active upon handling as untreated control mice |
| 3 | Exhibit a slower response to a tap on the cage and that were passive or docile when handled but still curious when alone in a new setting |
| 4 | Exhibit a lack of curiosity and little response to stimuli and that appear quite immobile |
| 5 | Exhibit labored breathing and are unable or slow to right themselves after being rolled onto their backs (moribund) |
| 6 | Dead mouse |
Supplementary Table 1: Scoring system for mice infected with influenza (1-6)
| Description | Score |
| --- | --- |
| Peri-vascular infiltrate​ | 0-3​ |
| Peri-bronchial infiltrate​ | 0-3​ |
| Epithelial damage​ | 0-2​ |
| Alveolar inflammatory exudate​ | 0-2​ |
Supplementary Table 2: Scoring system for Day 8 p.i. histology

### Slide 2
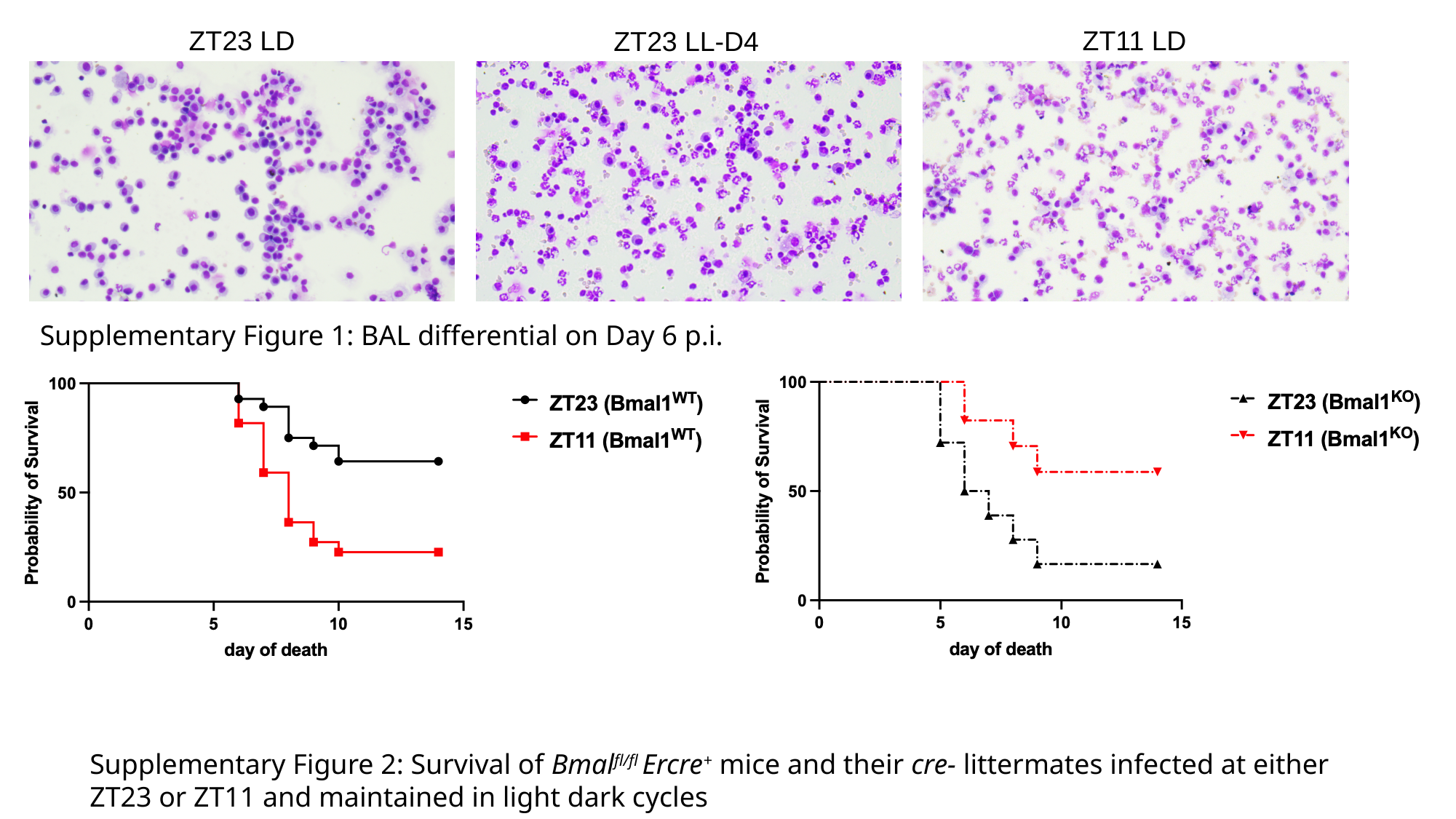

ZT11 LD
ZT23 LD
ZT23 LL-D4
Supplementary Figure 1: BAL differential on Day 6 p.i.
Supplementary Figure 2: Survival of Bmal^fl/fl^ Ercre^+^ mice and their cre- littermates infected at either ZT23 or ZT11 and maintained in light dark cycles

### Slide 3
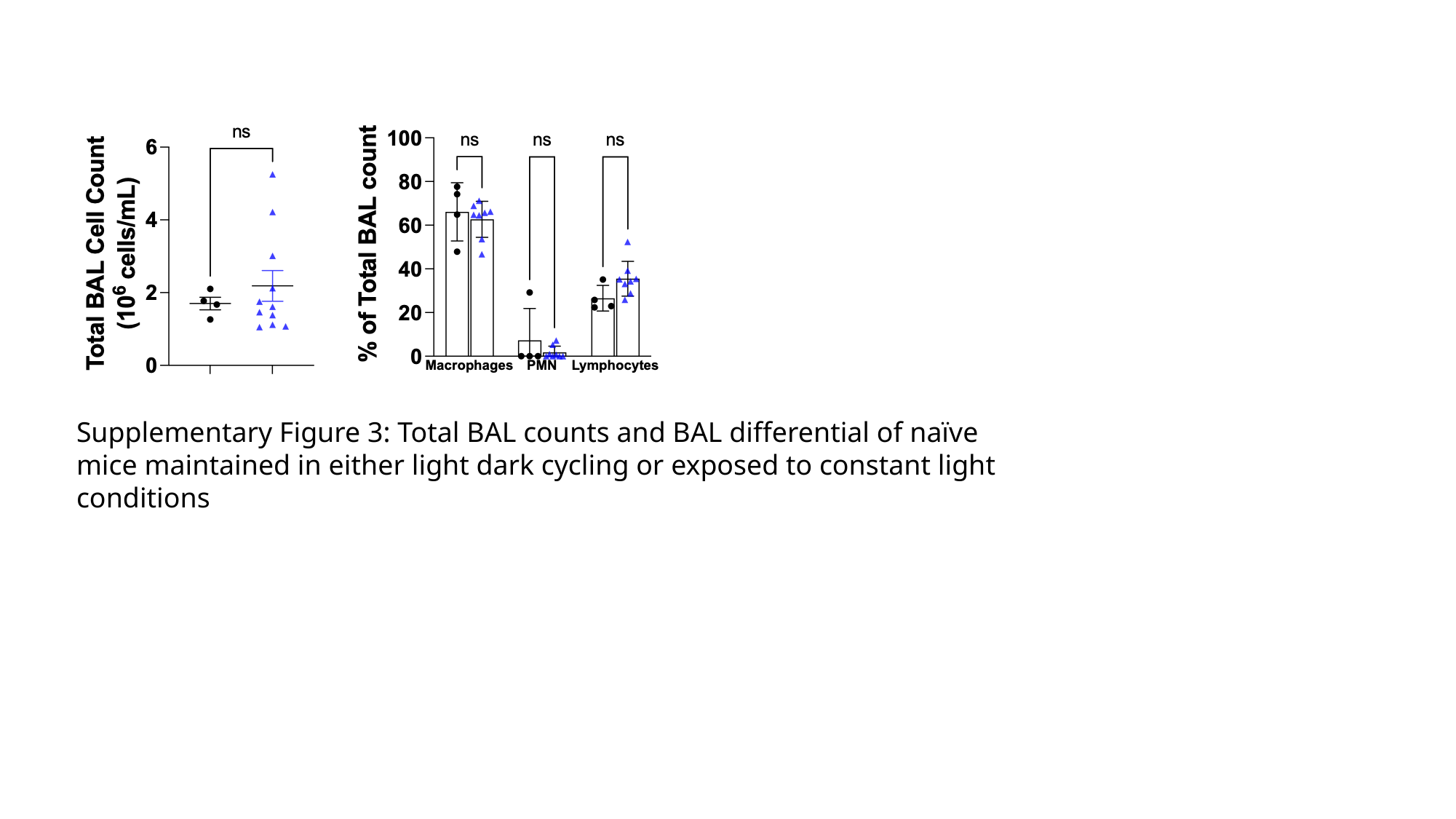

Supplementary Figure 3: Total BAL counts and BAL differential of naïve mice maintained in either light dark cycling or exposed to constant light conditions
